# Biophysical Modeling of Thalamocortical Circuit Dynamics: Species-Specific Insights into Neural Synchrony, Sleep Spindles, and Mechanisms of Neuropsychiatric Disorders

**DOI:** 10.64898/2026.02.01.703170

**Authors:** Basilis Zikopoulos, Natalia Matuk, Irina Romanova, Arash Yazdanbakhsh

## Abstract

Thalamocortical circuits play a fundamental role in cognitive functions, and neural synchronization, with disruptions implicated in disorders. Here, we investigated the neural dynamics of thalamocortical connectivity using computational modeling of rodent and primate thalamocortical loops. We incorporated distinct projections and varying network configurations and examined their impact on circuit synchrony, spiking patterns, and sleep spindle generation. Circuits included distinct core and matrix thalamocortical projections, with core pathways providing focal, driving input to middle cortical layers, while matrix pathways mediate widespread, modulatory signaling across superficial layers, and the presence of thalamic interneurons, which are scarce in rodents but comprise up to a third of the thalamic neurons in primates. In our simulations, these distinctions produced clear species-and loop architecture-dependent effects: rodent circuits were markedly more sensitive to parameter changes in core and matrix thalamocortical connectivity strength, while primate circuits maintained relatively stable spatiotemporal patterns across parameter variations, exhibiting greater stability and synchrony. Sleep spindle analysis likewise revealed species differences. Overall, across all thalamocortical configurations, rodent simulations produced spindles with greater spatiotemporal variability, showing irregular event structure and timing. In contrast, primate spindles were more uniform and coherent, with clearer and more consistent organization across neurons and time. These findings provide insights into species-specific differences in thalamocortical dynamics and have implications for modeling sensory and cognitive disruptions in disorders such as autism and schizophrenia. By incorporating distinct configurations, and interspecies differences, our model contributes to understanding how thalamocortical dysregulation may differentially impact spindle generation, network synchrony, and information processing across species.

## Introduction

The thalamocortical (TC) circuit, composed of the cortex, thalamus, and the inhibitory thalamic reticular nucleus (TRN), plays a critical role in both wake and sleep states (Zikopoulos & Barbas, 2007a). During wakefulness, the TC circuit is involved in sensory processing and attention (Zikopoulos & Barbas, 2007a; Whyte et al., 2024; Pinault, 2004), while during sleep, it generates sleep spindles – brief oscillations in the 10-16 Hz range commonly detected via electroencephalogram (EEG) recordings – and facilitates memory consolidation (Latchoumane et al., 2017; Manoach & Stickgold, 2019a). These oscillations have also been implicated in neurodevelopmental disorders such as schizophrenia and autism (Castelnovo et al., 2025; Gerardo & Manuel, 2020; Manoach & Stickgold, 2019b; Mylonas et al., 2022).

Two parallel and distinct sub-circuits form the basic organizational units of TC circuits: the core and the matrix. Core circuits are involved with sensory and cognitive processing and consist of focal projections, whereas matrix circuits are involved with limbic processing and memory consolidation, consisting of diffuse and widespread projections. Although both sub-circuits are found in all mammals, key differences exist between species. The significant expansion and specialization of the primate thalamus and cortical areas that have no homologues in rodents (Arcaro et al., 2015; Chartrand et al., 2023; García-Cabezas et al., 2022a; Jorstad et al., 2023; Joyce et al., 2022; Kim et al., 2023; Mengxing et al., 2023; Saalmann et al., 2012; Timbie et al., 2020), results in a clearer separation of core and matrix TC circuits, compared to rodents that differ by laminar distribution of cortical and thalamic projection neurons and their termination patterns, and neurochemical profiles in thalamus (Jones, 2007; Murray et al., 2007; Xiao et al., 2009; Zikopoulos & Barbas, 2007b). Moreover, rodents possess only a limited number of thalamic interneurons located in a few thalamic nuclei, while in primates, interneurons make up approximately a third of all thalamic neurons (Arcelli et al., 1997). This interspecies variability makes it crucial to examine thalamocortical dynamics across both rodents and primates. Another important distinction lies in the organization of TRN-thalamic interactions. These can be arranged in closed-loop (reciprocal innervation between TRN and the same thalamic neurons they inhibit), open-loop (no such reciprocity), or hybrid configurations. Despite extensive research on the role of the TC circuit in awake processing and sleep spindle generation, the specific contributions of core versus matrix circuits, TRN-thalamic connectivity configurations, and local inhibition, as well as how these lead to functional differences between species remain less understood.

To address this gap, we developed a detailed biophysical computational model of the rodent and primate thalamocortical systems, incorporating both structural and functional species-specific properties, including core, matrix, and mixed circuits, with the mixed circuit representing a functional combination of the two via cortico-cortico connection. Each species model was simulated across closed, open, and hybrid loop configurations. The hybrid loop represents a distinct loop configuration that integrates the features of both open and closed architectures. The model incorporated distinct and opposite receptor dynamics for each circuit: in the core, ionotropic receptors mediated thalamocortical connections and metabotropic receptors mediated corticothalamic connections, whereas in the matrix the arrangement was reversed. We controlled the spatial spread of neural activity using a difference-of-Gaussians convolution: the core used a narrow filter to restrict projections, while the matrix employed a wider filter for diffuse projections. Activity was initiated by external input to a single thalamic or TRN neuron, inducing TRN bursting and subsequent rebound spiking in thalamocortical neurons. This interaction initiated spindling, which we characterized by averaging multi-neuron activity and applying a 9–13 Hz bandpass filter to extract spindle signals.

Our findings highlighted faster synchronization with higher spindle density and amplitude in rodents compared to primates and pointed to mechanisms for differential spindle vulnerability in disorders; core spindles were more vulnerable to thalamocortical manipulations overall, but matrix manipulations were more effective at reducing overall spindle densities.

## Methods

### Thalamocortical Anatomy & Connectivity

The thalamocortical model includes cortical (Ctx), thalamocortical relay (TC), and thalamic reticular nucleus (TRN) neurons in the rodent and primate models; the latter also contains local thalamic inhibitory interneurons (IN). Core and matrix circuits interact at the cortical level, incorporating inter- and intra-cortical area mixing. Thalamic microarchitectural differences have been debated previously and implemented into this simulation (Brown et al., 2020; Willis et al., 2015; Yazdanbakhsh et al., 2023). Our model contains three different configurations: the closed, open and hybrid loop to understand how it impacts the activity of the circuit as a whole (Figure 1A-B). The closed loop refers to a configuration in which the TRN neuron inhibits the thalamic relay (TC) neuron, which reciprocally excites the original TRN neuron. In contrast, the open loop describes a configuration where the TRN neuron inhibits a TC neuron without reciprocal excitation. The hybrid loop combines features of both closed and open configurations (Yazdanbakhsh et al., 2023). Because empirical data on thalamoreticular connectivity remain limited, with existing anatomical studies suggesting that both open and closed motifs coexist within the thalamocortical system, a purely open or purely closed design would fail to capture the full range of biologically plausible dynamics. Incorporating a hybrid loop therefore provides an approximation of real-world circuitry, allowing us to simulate interactions that likely occur in vivo.

**Figure 1.**
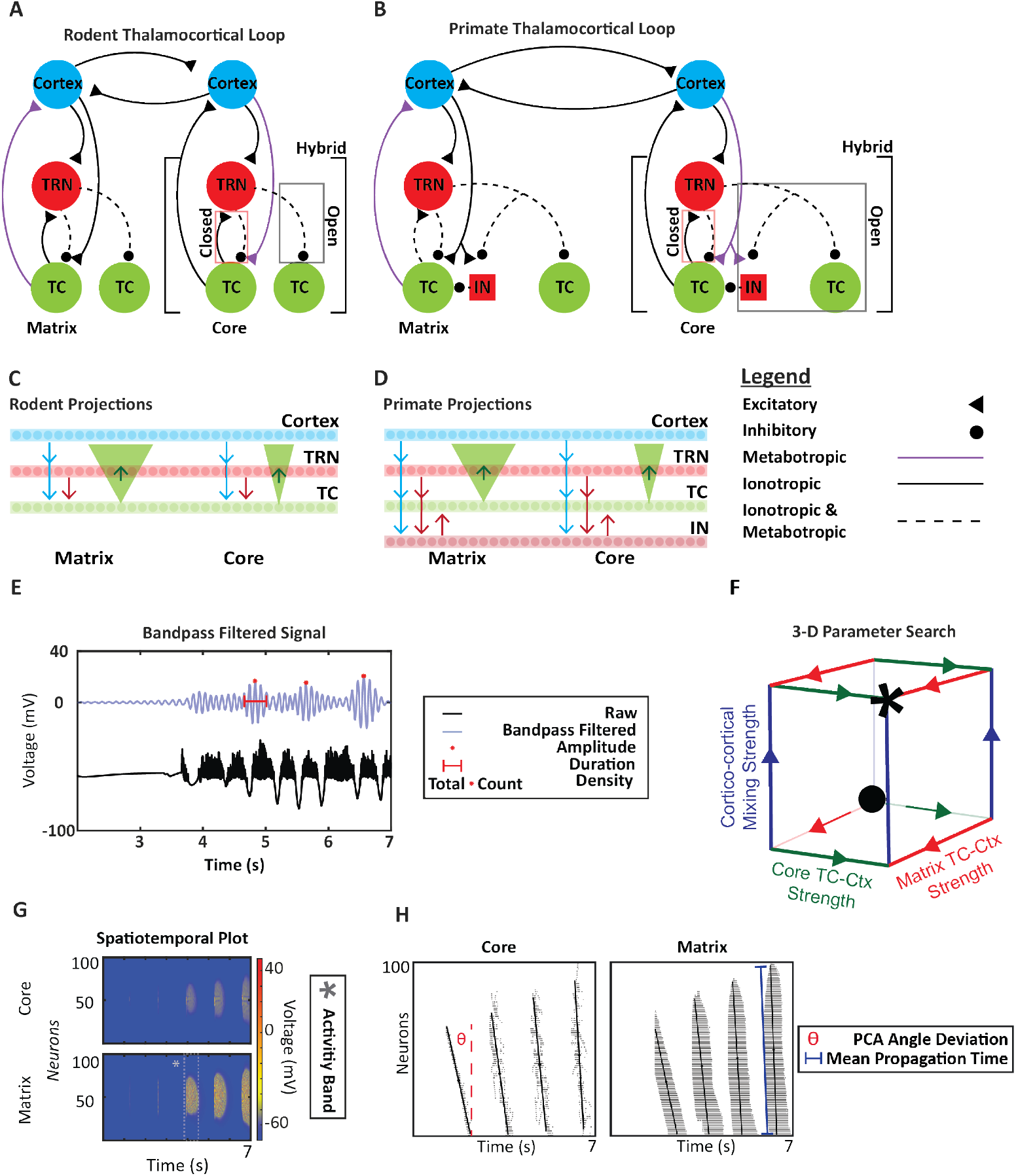
Experimental design and thalamocortical loop connectivity. **A**,**B.** The rodent and primate thalamocortical model. The cortical (blue), thalamocortical relay (green) and thalamic reticular nucleus (red circle) neurons make up the thalamocortical loop in rodents and primates. Local thalamic inhibitory interneurons (red square) are an additional component of the primate circuitry. The different types of synapses and receptors mediating the connections are labeled in the legend. GABA_A_ (ionotropic) or GABA_B_ (metabotropic) mediate inhibitory synapses. AMPA (ionotropic) or mGluR (metabotropic) mediate excitatory synapses. The core circuit labels microarchitectural differences (closed, open, and hybrid). The closed loop is defined as thalamic reticular neurons inhibiting thalamic relay cells that reciprocally excite them. The open loop is defined as thalamic reticular neurons inhibiting thalamic relay cells that do not reciprocally excite them. The hybrid loop is defined as a combination of the open and closed loop. **C, D**. Rodent and Primate Projections. All connections were one to one except for thalamocortical projections. The matrix thalamocortical projections were an order of magnitude wider than the core thalamocortical projections through the use of Gaussian filters and convolution. **E**. Sleep Spindle Detection. The raw signal is the averaged neural activity, while the blue is the bandpass-filtered signal with the frequency range of 9 –13 Hz. The attributes of spindles that were of interest are present in the legend. **F**. The 3-D cube template represents the three-dimensional parameter search of core and matrix thalamocortical mixing strength and cortico-cortical mixing strength. The * indicates the control case and the • indicates the starting case. **G**. Sample spatiotemporal plot of core and matrix cortical activity. The dashed gray box highlights an activity band. **H**. Raster plot of core and matrix cortical activity, with a PCA-derived eigenvector is overlaid on each raster plot. Its deviation from vertical (θ) measures synchrony, and the distance between its endpoints is used to compute the mean propagation time (MPT), quantifying temporal propagation across neurons.

Core and matrix circuits differ in their neurochemical and neuroanatomical properties. Based on anatomical and physiological findings (Reichova & Sherman, 2004; Zikopoulos & Barbas, 2007b), NR1 ionotropic receptors mediate matrix CTX-TC synapse, while mGluR1a metabotropic receptors mediate core CTX-TC synapse. These receptor type differences were implemented in our model (Figure 1C-D). For the reciprocal thalamocortical connections, receptor types were reversed to maintain their driving/modulatory roles of the core and matrix pathways. Furthermore, the core and matrix thalamocortical projections differ in their target regions and spread. Matrix thalamocortical projections target the superficial cortical layers in a widespread manner, while core thalamocortical projections target the middle/deep cortical layers more focally (Jones, 1998; Zikopoulos & Barbas, 2007b). Gaussian filters and convolution were used to simulate these differences, with matrix projections having a spread an order of magnitude wider than core projections.

### Network Geometry

The rodent model had a total of 600 neurons, while the primate had 800 neurons, with 100 neurons allocated to each neural type. Relative neuron count influence was approximated by scaling synaptic conductances to reflect differences in effective connectivity between neural populations. All neurons were modeled as single-compartment units governed by first-order differential equations that incorporate voltage-gated, intrinsic and synaptic currents. The simulation activity was initiated through a brief (50 ms) pulse to the circuit, and each simulation of the model ran for seven seconds.

### Voltage-Gated, Intrinsic and Synaptic Currents

The dynamics of each neuron are described by first-order differential equations based on the original work by Hodgkin and Huxley (1952), with adaptations from Pospischil et al. (2008) to capture additional physiological features. Each neuron incorporates Hodgkin-Huxley equations to describe sodium (I_Na_), potassium (I_K_) and leak (I_L_) currents, along with intrinsic currents specific to the physiological properties of the neural type and the synaptic currents. All currents were modeled based on Hodgkin-Huxley kinetics.

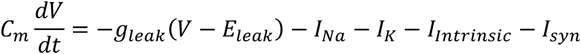

Each voltage-gated and intrinsic current has the following format:

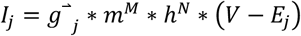

(Pospischil et al., 2008)

The maximal conductance of each current is denoted as *g*_*j*_. The gating variables *m* and *h* are probabilistic values that control the activation and deactivation kinetics, respectively. The driving force for the current is expressed as (*V – E*_*j*_), where *E*_*j*_ is the reversal potential.

### Hodgkin-Huxley Currents

The Hodgkin-Huxley consists of three primary currents: the voltage-dependent Na^2+^ current, the voltage-dependent K^+^ “delayed-rectifier” current, and the leakage current (Hodgkin & Huxley, 1952). The equations for all three currents are provided below and all necessary functions and constants are provided in Table 1 and Table 2.

### Sodium Current

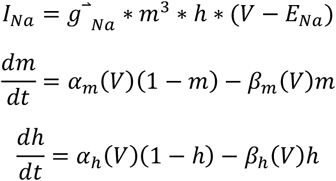

(Pospischil et al., 2008)

*Potassium “Delayed-Rectifier” Current:*

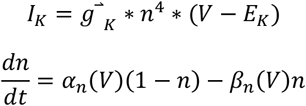

(Pospischil et al., 2008)

### Leakage Current

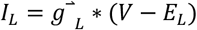

### Intrinsic Currents

To capture the distinct physiological properties of TRN and TC relay neurons, two additional currents were incorporated into the model. Both neuron types possess a low-threshold Ca^2+^ current (I_T_) that is responsible for their bursting and rebound bursting properties; however, the kinetics of this current differ between the TRN and TC neurons, thus requiring distinct equations (Bal & McCormick, 1993; A. Destexhe et al., 1993; Huguenard & Prince, 1992; McCormick & Huguenard, 1992). In addition, TC relay neurons possess a hyperpolarization-activated current (I_H_), which facilitates the repolarization of the TC membrane potential following hyperpolarization (A. Destexhe et al., 1996; Huguenard & Prince, 1992). All constants and necessary functions for each current are listed in Tables 1 and Table 2.

### Thalamic Reticular Nucleus T-Current

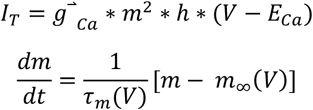

(A. Destexhe et al., 1994)

### Thalamocortical Relay Neuron T-current

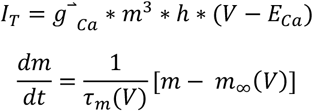

(A. Destexhe et al., 1993)

### Thalamocortical Relay Neuron H-current

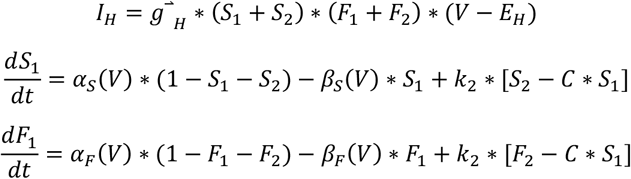

(A. Destexhe et al., 1993)

### Synaptic Currents

Synaptic communication in the model incorporates the following neurotransmitter receptors: AMPA, NMDA, GABA_A_, GABA_B_, and mGluR. These ionotropic and metabotropic receptors mediate the synaptic connections.

### Ionotropic Channels

The current for ionotropic receptors is modeled using the following equation:

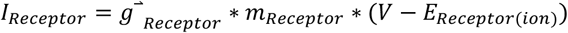

The maximal conductance, g_receptor_, of the ion associated with the specific receptor; the total fraction of open post-synaptic receptors, m_receptor_, and the driving force (V-E_receptor (ion)_) control the flow of the ions through the specific receptor. The fraction of open receptors is dependent on the concentration of neurotransmitter present. The generic differential equation describing this relationship is as follows:

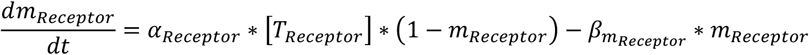

The parameters *α* and *β* represent the forward and backward binding rates, respectively, while [T_receptor_] represents the concentration of neurotransmitter present. All of the constants associated with each ionotropic receptor are present in Table 3.

### Metabotropic Channels

The format for the metabotropic receptor and all necessary equations are provided below (Destexhe et al., 1998). All the parameter values are provided in Table 4. The general current equation is provided below:

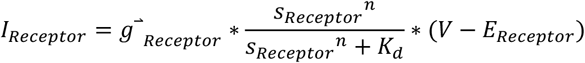

The variable,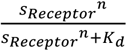, represents the concentration of second-messengers. The equation for the rate of change of the openness of the post-synaptic receptors is as follows:

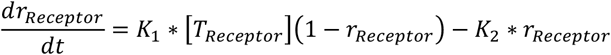

The variables, K_1_ = 1.3, and K_2_ = 0.006, are constant values similar to the forward and binding rates. An additional third differential equation describes the receptor desensitization rate:

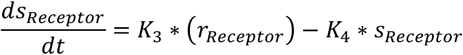

The sensitization and desensitization rates are represented by the variables K_3_ = 0.09, and K_4_ = 0.0064. The variable, *s*_*Receptor*_, represents second-messenger concentration that both contributes to the opening of the specific metabotropic receptors and to the decay of second-messenger concentration.

The metabotropic glutamate receptor (mGluR), responsible for excitatory signaling, was modeled using the kinetics and kinematics of the GABA_B_ receptor. The reversal potential for mGluR (E_mGluR_) was set to 0 mV, consistent with E_AMPA_.

### Thalamocortical Activity Synchronization & Statistical Analysis

The synchronization of thalamocortical activity was assessed both qualitatively and quantitatively. Three-dimensional parameter searches were conducted and outlined within a 3-D cube framework (Figure 1F). The x- and y-axes represent the relative prevalence (strength) of core and matrix thalamocortical connectivity, respectively, while the z-axis represents the strength of cortico-cortical mixing (Figure 1F). The starting condition of each thalamocortical loop iteration (•, Figure 1F) corresponds to the initial state of the loop, where all parameters are set to relatively low values. The control case of the network was defined as a balanced thalamocortical activity level, situated between complete inactivity and overly saturated activity, allowing for future investigation of neurobiological parameters.

At each vertex of the cube, spatiotemporal maps for the core and matrix thalamocortical activity were displayed, providing insights into the neural activity of a region (TRN, TC, IN and Ctx) over time (Figure 1G). To quantify the spatiotemporal structure of thalamocortical activity bands, we applied principal component analysis (PCA) to the spike-time raster data for each activity band. For each activity band, we z-scored the time and neuron axes to maintain a shared reference frame across species, loop configurations, and trials. PCA was then performed on the z-scored time–neuron activity band to obtain the first principal component (PC1), which reflects the orientation of the largest standard deviation of the activity bands and therefore indicate the orientation of the activity band in the spaciotemporal space. The PCA orientation angle (θ) was computed by projecting the PC1 loading vector onto the raster’s time-neuron plane and measuring its angle relative to vertical. Angles (θ) near 90° indicate vertically aligned bands in which neurons fire with minimal temporal offset (high synchrony). Angles (θ) deviating from 90° indicate tilted, propagating activity in which spiking progresses across neurons over time (lower synchrony).

During analysis of the closed-loop rodent model, we observed that most activity bands were bilaterally symmetric about the vertical axis and formed boomerang-shaped patterns. Because PCA extracts the axis of maximum variance, applying PCA to this fully symmetric structure produced an artificially vertical eigenvector, even when the underlying spiking pattern itself was not synchronous. The vertical eigenvector was therefore a geometric artifact of bilateral symmetry, not a reflection of true synchrony. To correct for this artifact and recover the true temporal structure of rodent closed-loop activity bands, we performed PCA on the upper half of each activity band only. Removing the symmetric lower half prevents symmetry from forcing PCA-derived eigenvector to align vertically. In this half-band representation, the orientation of the PCA-derived eigenvector reflects the actual spatiotemporal dynamics of the activity band, rather than the imposed geometry of bilateral symmetry. This procedure enabled valid comparisons across rastergrams of rodent and primate activity, and across closed, open, and hybrid loops. The same global z-scoring and angle-based synchrony measures were then applied uniformly across rastergrams.

We further used the PCA first principal component eigenvector orientation to compute the mean propagation time (MPT), defined as the elapsed time between the first and last spiking neurons within an activity band divided by the number of active neurons in that interval. To obtain this value, we embedded the PCA-derived eigenvector as it is stretched across the activity band, allowing us to estimate both the time spanned along this vector with the number of neurons it traversed. MPT therefore measures time offset per neuron in the sequence. Synchronous activity produces MPT values near zero, whereas propagating traveling wave-like activity yields larger MPT values reflecting lower activity band synchrony. PCA-derived orientation angles and MPT were jointly used to characterize the temporal structure of each activity band.

### Sleep Spindle Detection & Statistical Analysis

All simulations were conducted with core and matrix cortical mixing, meaning that core and matrix circuits interacted at the level of the cortex, through connections (mixing/integration) across layers or across areas. Sleep spindles for core and matrix activity were calculated by averaging the neural activity of five core and five matrix neurons at a time, separately from the functionally mixed thalamocortical circuit. Mixed spindles were calculated by averaging the combined activity of 10 cortical neurons (5 matrix and 5 core) from the functionally mixed thalamocortical loop. All averaged activity was bandpass filtered in the 9–13 Hz range (Figure 1E) for sleep spindle detection.

We assessed sleep spindle amplitude, duration, and density following the criteria outlined by Gonzalez et al. (2022). Spindles were considered for analysis, if their amplitude exceeded a set threshold (30% of peak amplitude spindle) and was not within 0.5 seconds of another detected amplitude (Gonzalez et al., 2022). Spindle duration was defined as the time difference between two local minima in reference to a detected amplitude. Spindle density was calculated by counting the number of amplitudes within a bandpass-filtered averaged activity array. All spindle metrics were visually cross-validated to ensure accuracy.

Because spindle metrics (amplitude, duration and density), exhibited non-normal distributions, we performed rank-ordered statistical analysis. Kruskal-Wallis tests were performed to evaluate statistical significance among rodent and primate spindle metrics (sleep spindle amplitude, duration and density) across all three loop configurations (closed, open, hybrid).

We computed the fast Fourier transform (FFT) of averaged time-domain signals to analyze spectral power in rodent and primate thalamocortical circuits across open, closed, and hybrid loop configurations. Power spectral densities (PSDs) were then plotted on a logarithmic scale within the 8–20 Hz range to capture both prominent spindle peaks and subtle spindle-related dynamics. Similarly to the approach of Fernandez & Lüthi (2020), our goal was to continuously assess thalamocortical spindle activity rather than focus solely on discrete spindle detection. By examining the spectral distribution, we aimed to detect species- and condition-specific differences in content and organization of frequencies within the spindle range, including broad sigma-band shoulders and distributed frequency components. This spectral approach is particularly relevant for comparing rodent and primate circuits, where differences in spindle expression may reflect species-specific adaptations in thalamocortical architecture.

## Results

### Activity in Rodent and Primate TC Circuit Across the Parameter Search

In both species, the closed loop produced symmetrical, vertically aligned activity bands (Figure 2). In contrast, the open and, to a lesser extent, hybrid loops generated asymmetrical bands characteristic of traveling waves (Figures 3–4). Traveling waves were most prominent in the rodent model but were also observed under certain parameter settings in the primate model.

**Figure 2.**
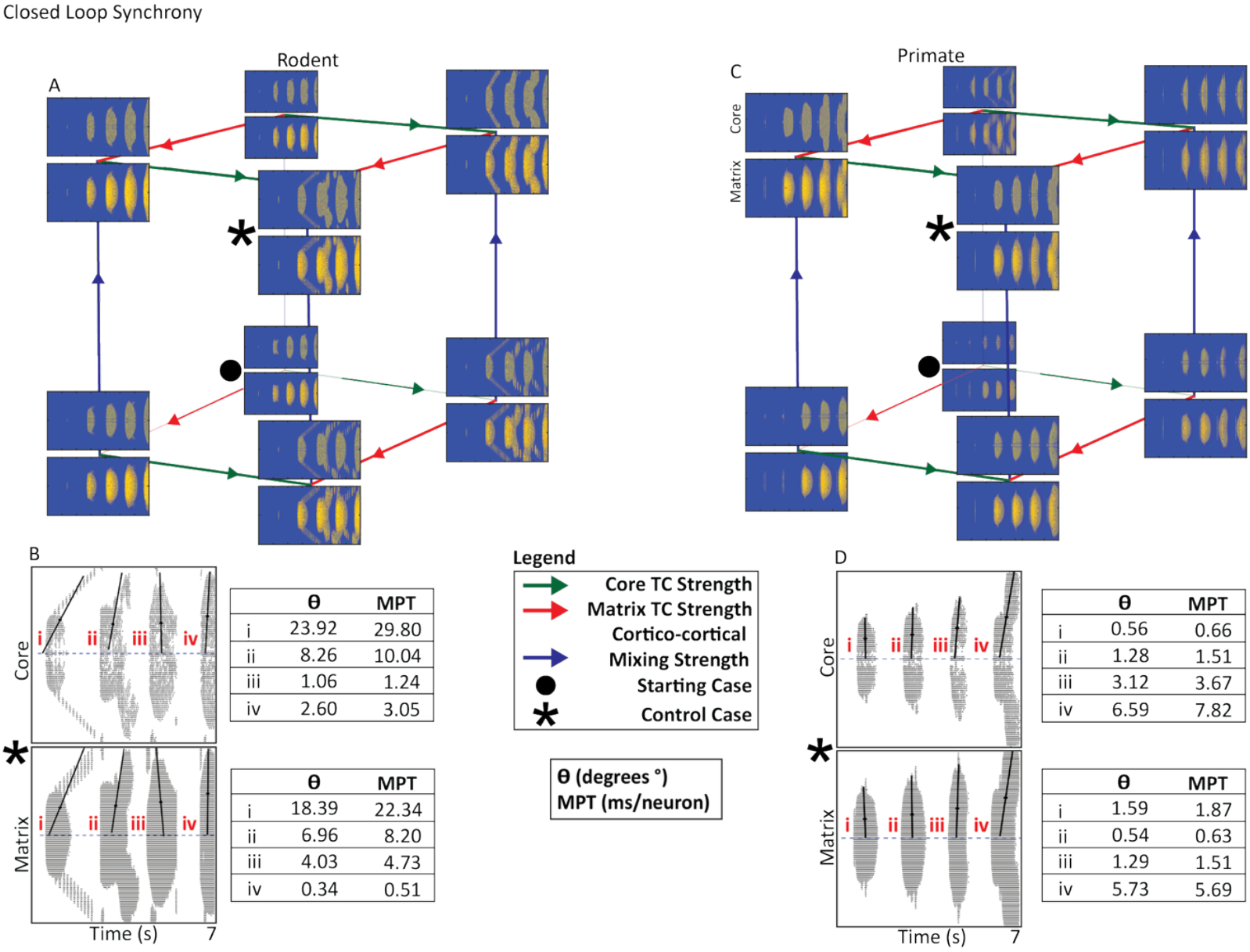
Closed Thalamocortical Loop Cortical Synchrony. **A, C.** Rodent and primate cubes present the three-way parameter search of core and matrix thalamocortical strength as well as cortico-cortical mixing strength. The • indicates the starting case, while the * indicates the control case. We assessed changes in synchrony using the control case. Primate cortical activity synchrony and spread was higher than rodent cortical activity. **B, D**. Raster plots present control case rodent and primate cortical activity from the core and matrix. Each band of activity analyzed is marked with a roman numeral. Propagation angle (θ) and mean propagation time (MPT) are shown for each activity band. θ reflects burst synchrony based on the orientation of the PC1 eigenvector, while MPT was derived from the slope of this eigenvector, representing temporal offset per neuron during burst propagation. Larger θ and MPT values indicate slower, more sequential propagation, whereas smaller values reflect more synchronous activation (see *Thalamocortical Activity Synchronization & Statistical Analysis* section).

**Figure 3.**
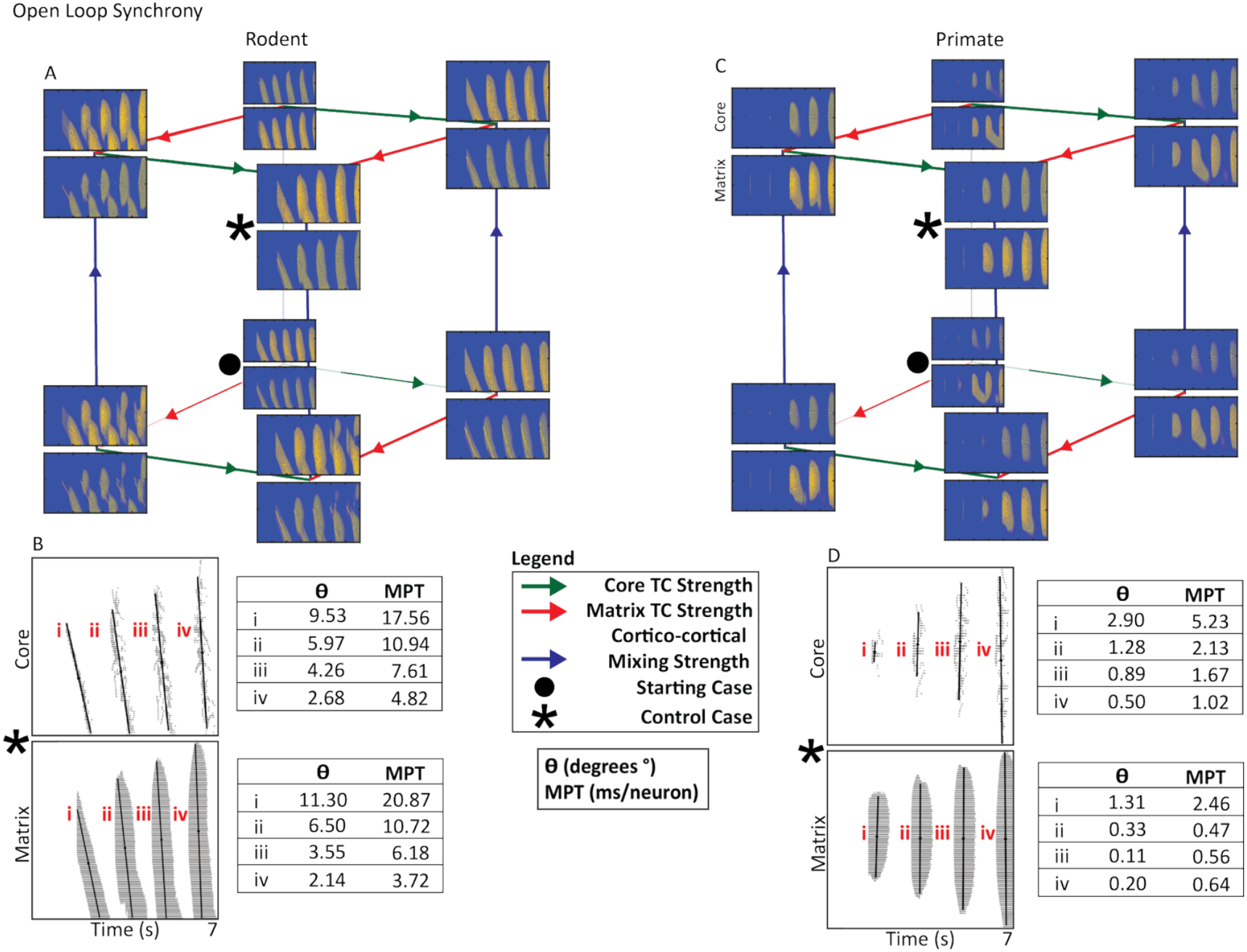
Open Thalamocortical Loop Cortical Synchrony. **A, C.** Rodent and primate cubes present the three-way parameter search of core and matrix thalamocortical strength as well as cortico-cortical mixing strength. The • indicates the starting case, while the * indicates the control case. We assessed changes in synchrony using the control case. Primate cortical activity synchrony and spread was higher than rodent cortical activity. Rodent cortical activity contained higher amounts of traveling waves that are viewed as inherently asynchronous. **B, D**. Raster plots present control case rodent and primate cortical activity from the core and matrix. Each band of activity analyzed is marked with a roman numeral. Rodent activity bands encompassed more neurons than primate activity bands. Propagation angle (θ) and mean propagation time (MPT) are shown for each activity band. θ reflects burst synchrony based on the orientation of the PC1 eigenvector, while MPT was derived from the slope of this eigenvector, representing temporal offset per neuron during burst propagation. Larger θ and MPT values indicate slower, more sequential propagation, whereas smaller values reflect more synchronous activation *(see Thalamocortical Activity Synchronization & Statistical Analysis* section).

In both rodent and primate closed loops, the spread of cortical activity increased as a function of core and matrix thalamocortical strength. In the rodent model, increasing core thalamocortical strength altered the synchrony of cortical activity, producing more tilted activity bands and traveling-wave structure, whereas the primate model was largely unaffected. Additionally, increased cortico-cortical mixing amplified the spread of cortical activity in both rodent and primate closed loops (Figure 2A & 2C). Differences in the effect of core thalamocortical strength on synchrony led to drastically different control cases in the rodent and primate.

The rodent open loop generated traveling waves, regardless of the values of core TC strength, matrix TC strength, or cortico-cortical mixing. The primate open loop, while still capable of generating asymmetrical bands in some parameterizations, often produced more organized, less prominent traveling waves, partially resembling the closed loop geometry (Figure 3C •). Across the parameter search, neither model showed major changes in the spread of activity as a function of increased core or matrix thalamocortical strength. In the rodent open loop, increasing matrix thalamocortical strength caused separation in the traveling waves (Figure 3A). In the primate open loop, increased matrix, and to a lesser extent, core thalamocortical strength, reduced the prominence of traveling waves (Figure 3D).

In the rodent hybrid loop, increasing core TC strength resulted in more synchronized and cohesive activity bands (Figure 4A & 4C). In contrast, increasing matrix TC strength heightened traveling waves and separation. For the primate hybrid loop, increasing cortico-cortical strength caused an increase in the spread of cortical activity. However, even at the most extreme parameters, primate activity bands remained substantially more synchronous compared to rodents (Figure 4B & 4D).

**Figure 4.**
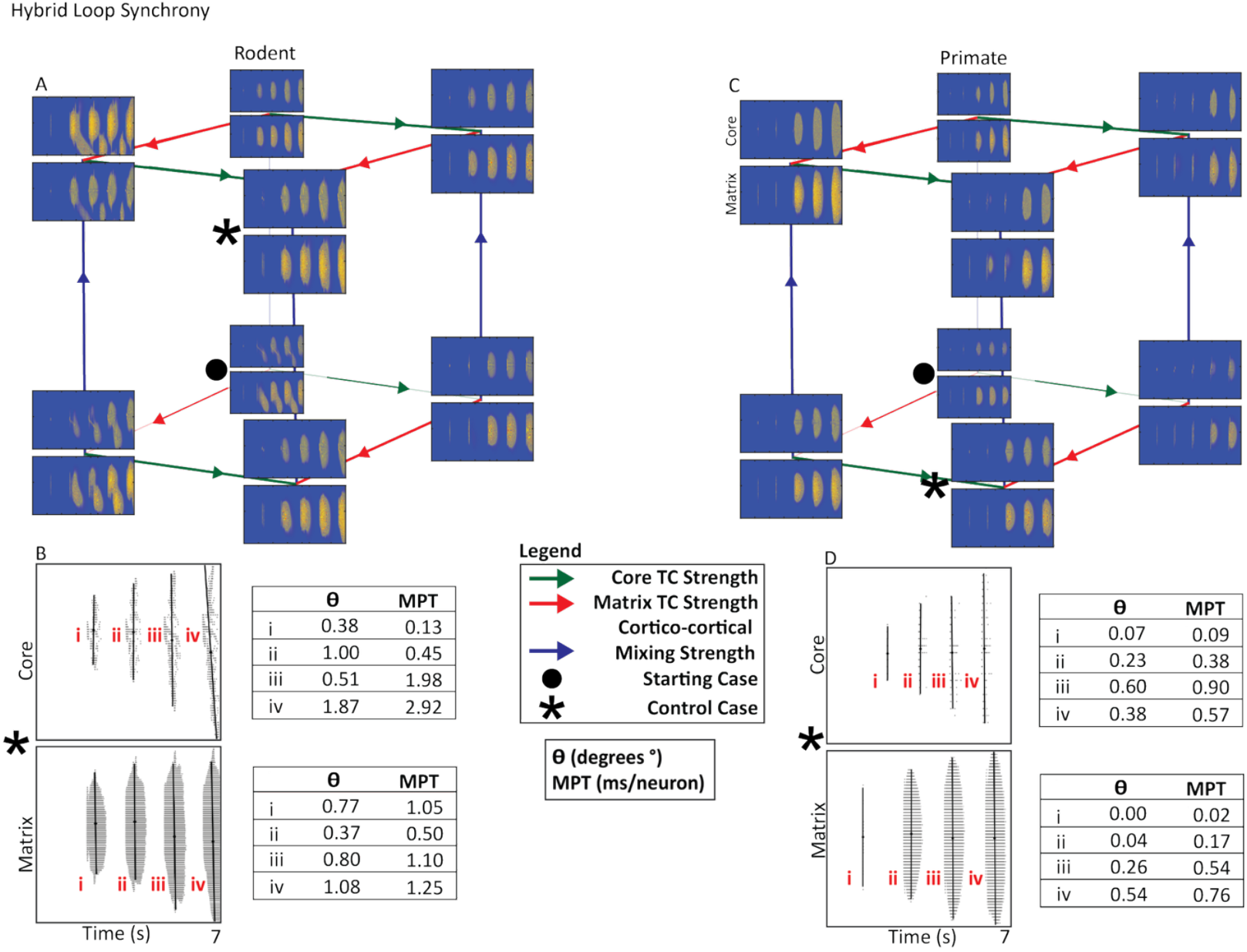
Hybrid Thalamocortical Loop Cortical Synchrony. **A, C.** Rodent and primate cubes present the three-way parameter search of core and matrix thalamocortical strength as well as cortico-cortical mixing strength. The • indicates the starting case, while the * indicates the control case. The control case determined in the hybrid loop required lower levels of cortico-cortical mixing. We assessed changes in synchrony using the control case. Primate cortical activity was more synchronized than rodent cortical activity. **B, D**. Raster plots present control case rodent and primate cortical activity from the core and matrix. Each band of activity analyzed is marked with a roman numeral. Propagation angle (θ) and mean propagation time (MPT) are shown for each activity band. θ reflects burst synchrony based on the orientation of the PC1 eigenvector, while MPT was derived from the slope of this eigenvector, representing temporal offset per neuron during burst propagation. Larger θ and MPT values indicate slower, more sequential propagation, whereas smaller values reflect more synchronous activation (see *Thalamocortical Activity Synchronization & Statistical Analysis* section).

Before going into the key findings from each loop configuration, we wanted to note general trends that were present in all three versions of the thalamocortical loop and in both species. Matrix cortical activity tended to synchronize more than the core cortical activity. Across the parameter search for both species, matrix-derived cortical activity appeared more diffuse than core activity, although this difference was far more pronounced in rodents. The cortex exhibited the highest synchrony compared to the TRN, IN (in primates), and TC relay neurons. TC relay neurons consistently demonstrated the synchrony across the thalamocortical loop.

### Spatiotemporal Structure of Thalamocortical Activity Across Loop Configurations

Across simulations, thalamocortical dynamics showed configuration- and species-dependent differences in the temporal organization of activity bands. PCA-derived eigenvector angle and mean propagation time (MPT) revealed consistent patterns in the temporal structure of TC activity bands. Rodent network systematically produced activity bands with greater temporal dispersion, whereas primate networks generated more tight activity. These species differences were evident across open, closed, and hybrid loop configurations and were expressed similarly across core and matrix pathways.

### PCA- and MPT-Based Activity Band Structure and Dimensionality

Activity band angles in rodents tended to deviate more from the vertical, indicating that rodent activity bands exhibited stronger temporal shift and a greater degree of sequential activation across the neuronal population. Correspondingly, rodent MPT values were consistently higher, reflecting a larger elapsed time per neuron along the activity band. Together, these metrics showed that rodent TC circuits tended to exhibit activity that travels gradually across neighboring neurons rather than occur simultaneously. Primate loop configurations showed the opposite pattern, with near vertical PCA orientations, indicating minimal temporal tilt and a high degree of activity concurrence within each band. Primate MPT values were accordingly small, indicating minimal temporal spread across neurons. These results may suggest that primate TC loops support more synchronized population events.

Loop configurations further regulated these species-specific tendencies. Open and closed-core circuits amplified the differences, exhibiting the strongest propagation patterns in rodent and the most synchronous activity in primates. In contrast, hybrid configurations were highly synchronous in both species. In these hybrid loops, PCA angles were consistently near vertical and MPT values were also small, indicating that the mixed connectivity pattern stabilizes synchronized firing regardless of species. Across all analyses, PCA and MPT showed consistent patterns: rodents tend to exhibit more sequential firing, while primate loops have more synchronized activity.

### Activity Synchrony of the Closed TRN-TC Loop

The primate closed loop consistently displayed higher levels of synchrony, regardless of cortico-cortical or core and matrix thalamocortical connectivity changes, unlike the rodent closed loop (Figure 2A,C). While both species exhibited symmetrical activity bands in the closed architecture, rodent synchrony level was more sensitive to parameter variation: increases in core TC strength altered the spatiotemporal pattern of rodent cortical activity, whereas primate activity retained comparatively stable synchrony level. These differences were also reflected in the PCA and MPT analyses (Figure 2B,D).

Rodent PCA angles deviated more prominently from vertical, notably in the core condition with a 23.92° deviation from 90°, corresponding to an MPT of 9.59 ms/neuron, indicating strong temporal tilt and sequential propagation. In contrast, primate closed-core PCA deviations clustered near vertical (0.56°– 6.59°) with smaller MPT values (1.38–3.98 ms/neuron), reflecting more synchronous activation. Similar patterns emerged in the matrix pathway: rodent deviations ranged 2.65°–6.22° (MPT: 0.45–4.54 ms/neuron), whereas primate deviations remained tighter (1.67°–3.13°) with lower MPTs (0.94–3.28 ms/neuron).

Additionally, we observed species-specific differences in how synchrony evolved across the activity bands over the time course of the simulation. In primates, core and matrix cortical bands began in a highly synchronous configuration and remained vertically aligned throughout the progression of the simulation, with PCA orientations remaining close to 90° and consistently minimal MPT values (Figure 2C,D). In contrast, rodents began in a less synchronized state, with tilted activity bands in the early stages of the simulation, which then became more vertically organized as the simulation progressed (Figure 2A,B). This gradual synchronization of rodent activity bands was reflected in PCA deviations that moved closer to 0° and in decreasing MPT values over the course of the simulation. Thus, while primate closed-loop synchrony was stable and appeared inherent, rodent synchrony likely had to develop over time.

### Activity Synchrony of the Open TRN-TC Loop

The rodent open loop was dominated by traveling waves regardless of the cortico-cortical, core or matrix thalamocortical connectivity (Figure 3A). PCA captured this pattern of sequential propagation: rodent activity band orientations showed clear deviations from vertical (2.14°–11.30°) along with elevated MPT values (3.72–20.87 ms/neuron) (Figure 3B). The primate open loop also exhibited traveling-wave activity, but these waves were less pronounced than those observed in rodents (Figure 3C). Accordingly, primate activity band deviations remained small (0.87°–2.90°) and produced substantially lower MPT values (0.47–5.23 ms/neuron) than rodents, indicating higher synchrony. In the primate open loop, increased core or matrix TC strength reduced the prominence of traveling waves, resulting in a more synchronous activity pattern (Figure 3D). In contrast, in the rodent open loop, increases in matrix TC strength caused greater separation among traveling waves (Figure 3A). As such, PCA and MPT jointly indicated higher synchrony in the primate compared to the rodent model.

### Synchrony Across Circuit Nodes in the Hybrid TRN-TC Loop

Hybrid-loop architecture produced the most synchronized patterns of activity bands for both species. Rodent hybrid activity was very sensitive to the parameter changes in connectivity, as increasing core TC strength or cortico-cortical mixing produced more vertically aligned activity bands (Figure 4A), reflecting a shift toward synchrony. This was supported by PCA deviations that approached zero 0.38°–1.87° in the core pathway; 0.77°–1.08° in the matrix pathway and MPT values were correspondingly small (Figure 4B), indicating reduced temporal dispersion across neurons. In contrast, increasing matrix TC strength had the opposite effect as activity bands became more asymmetric and more widely separated, producing traveling waves (Figure 4A). Primate hybrid loop remained more synchronous overall, even as core TC strength, matrix TC strength, and cortico-cortical mixing were varied (Figure 4C). Activity bands stayed nearly vertical, with PCA-derived eigenvector deviations consistently near zero (0.07°–0.60° in the core pathway; 0.00°–0.54° in the matrix pathway) and extremely small MPT values (0.02–0.90 ms/neuron) (Figure 4D), indicating near-synchronous population activation.

Together, the hybrid loop results reveal that rodent loops are capable of producing both synchronous and asynchronous activity, depending on whether core TC strength or cortico-cortical mixing dominates the circuit. By contrast, primate loops remain consistently synchronous.

### Sleep Spindle Metrics

We assessed sleep spindles across rodents and primates as a function of their reticulo-thalamic (closed, open or hybrid loop) connectivity. Figure 5A, B, and C, present examples of core and matrix sleep spindles generated by the three versions of the rodent and primate model. Across all three loop architectures, clear species differences were visible in the structure and consistency of spindle activity. In the rodent model, both core and matrix spindles appeared more spatiotemporally heterogeneous. The shaded activity bands varied noticeably in width and shape, and the number of neurons participating in each spindle shifted across events. In addition, the rodent model showed more pronounced traveling-wave structure and less uniform spacing of the activity bands from which spindle events were extracted.

**Figure 5.**
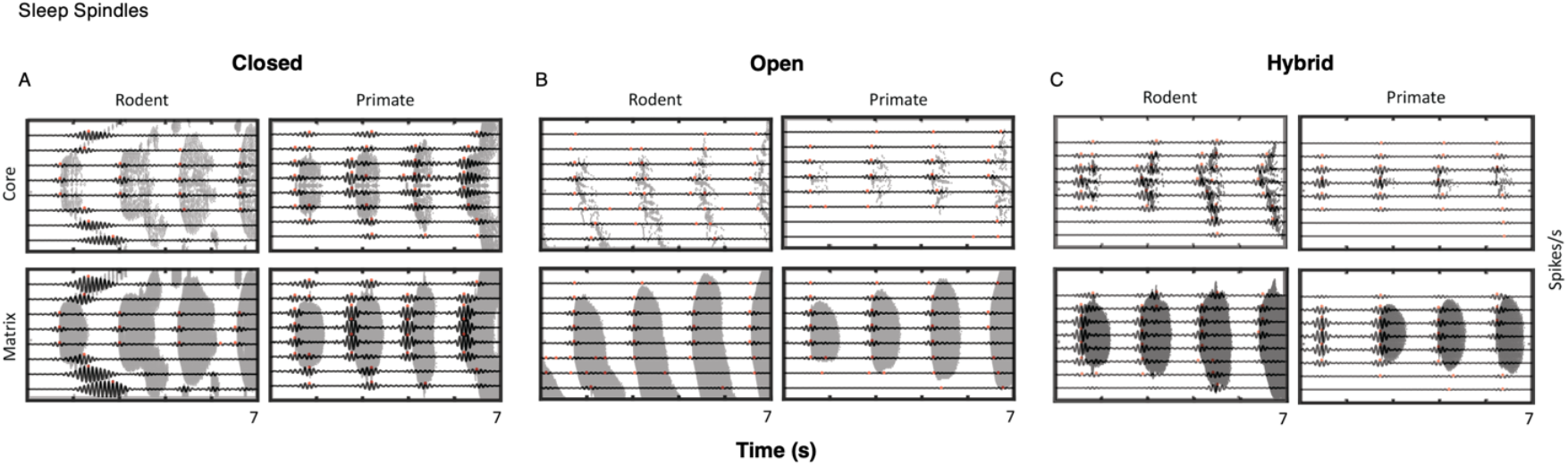
Rodent vs. Primate Thalamocortical Loop Spindles Across Loop Configurations. Core and matrix sleep spindle examples in rodent and primate models. **A, Closed Loop.** Rodent spindles appeared more variable and broader in shape in both core and matrix pathways, with no strong separation between the two. Primate spindles showed much more uniform structure across core and matrix, with clearer boundaries and more consistent timing across events. **B, Open Loop**. Rodent core and matrix spindles showed greater spatiotemporal variability, with spindle shapes that were less uniform and lower in amplitude, whereas primate spindles, both core and matrix, appeared to be more stable in shape and more regularly spaced. **C, Hybrid Loop**. Hybrid loop simulations showed the same overall pattern: there was no strong core-matrix distinction in either species, but rodent spindles were more variable overall, whereas primate spindles maintained a more uniform structure across events. Across all loop configurations, the rodent model displayed greater spindle spatiotemporal variability, while the primate

In the primate model, core and matrix spindles appeared more consistent and temporally aligned. The spindles were more regular in shape and had clearer boundaries. Successive spindles had more regular duration and spacing, and the overall pattern of activity appeared more stable across loop types. Primate core and matrix circuits displayed this increased regularity relative to the rodent model. Altogether, the spindle patterns indicate greater spatiotemporal variability in the rodent model, while the primate model shows more coherent and consistently structured spindles across core and matrix of all three thalamocortical loop configurations.

### Power Spectral Density Analysis of TC Activity

In rodents, the open loop condition for both core and matrix circuits (Figure 6C–D) showed elevated power extending into the 13–15 Hz range, accompanied by low-amplitude oscillations and scattered peaks between 11 and 13 Hz. In the closed loop, the matrix configuration (Figure 6H) exhibited a pronounced peak centered around 10–12 Hz, while the core configuration (Figure 6G) showed a smaller peak in the same range with continued elevated power into the 13–15 Hz band. In the hybrid loop (Figure 6K–L), persistent elevated activity between 13 and 15 Hz was observed in both core and matrix circuits, along with multiple smaller peaks between 11 and 14 Hz. Across all conditions, elevated power within the 13–15 Hz range was consistently present, though the spectral shape around 13–14 Hz tended to be flatter and less sharply defined than peaks at lower spindle frequencies.

**Figure 6.**
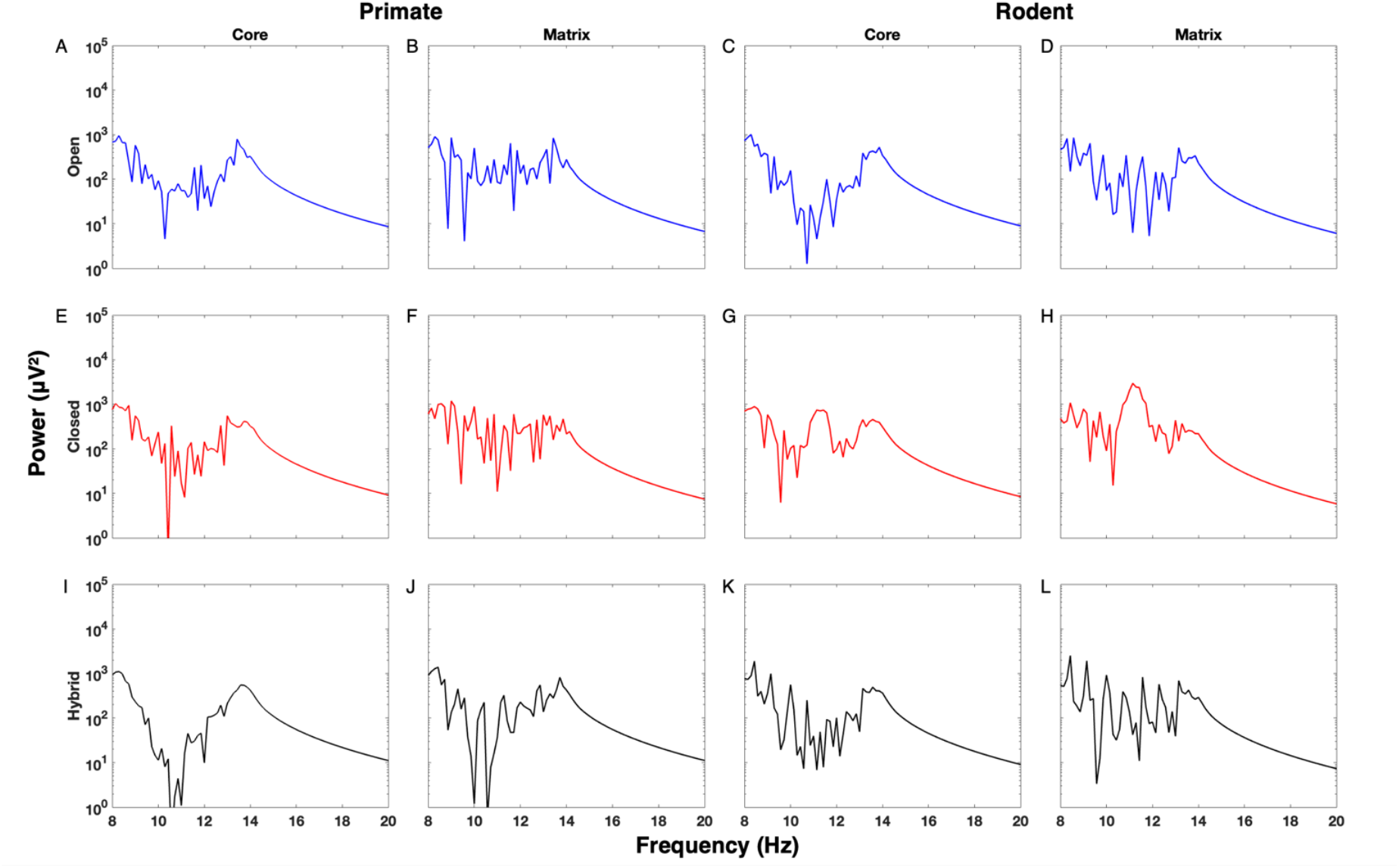
Spectral Power Across Rodent and Primate Thalamocortical Loop Configurations. Power spectral densities (PSDs) of averaged simulated signals were generated and shown here for thalamocortical core (left) and matrix (right) pathways under **open (A–D), closed (E–H)**, and **hybrid (I– L)** loop configurations in **rodent (A–B, E–F, I–J)** and **primate (C–D, G–H, K–L)** models. PSDs were plotted on a logarithmic scale from 8 to 20 Hz, covering the spindle frequency range (∼10–16 Hz) and adjacent bands. Open loop conditions (blue) showed low-amplitude, broad oscillations without distinct peaks. Closed loops (red) displayed sharper, higher-amplitude peaks centered in the spindle range. Hybrid loops (black) showed intermediate profiles, with partial peak formation and greater variability across core and matrix circuits. Primate PSDs were generally lower in amplitude and more broadly distributed than rodent PSDs.

In primates, the open loop configuration for core and matrix circuits (Figure 6A–B) showed persistent power in the 13–14 Hz range, with more scattered activity extending to 11 Hz. In the hybrid loop (Figure 6I–J), we observed a distinct peak between 13 and 15 Hz, while lower frequencies were less prominent. In the closed loop, only the core pathway (Figure 6E) showed slight increases in power around 13–14 Hz, though overall power did not exceed that of the open loop. The closed matrix configuration (Figure 6F) displayed a more uniform and flatter distribution compared to the core. Notably, in hybrid loops, especially in the core pathway (Figure 6I), a sharper and more defined peak emerged between 13 and 14 Hz, more distinctly expressed than in rodents at similar frequencies. Matrix activity remained relatively flat in both species, but primates exhibited slightly greater spindle-band power in the matrix pathway.

Both primate and rodent hybrid core circuits (Figure 6I,K) exhibited spindle power, though it was more pronounced in primates. In both species, matrix spindle activity was lower (Figure 6J,L), yet primates still showed slightly greater spindle prominence in the matrix compared to rodents. Overall, the rodent spectra displayed more diffuse and less sharply defined spindle-band structure across loop configurations, whereas primate spectra were more defined and showed consistent peaks. This suggests that the rodent model exhibits more variable and less rhythmic spindle activity compared to the primate model.

## Discussion

### Activity in Rodent vs Primate Thalamocortical Model Circuits

This study explored how several critical TC circuit organizational principles can affect thalamocortical network dynamics, in particular sleep spindle activity characteristics, with possible implications for future experimental work and computational modeling. Our findings revealed species-specific patterns of activity between distinct core and matrix thalamocortical circuits, each including three different thalamoreticular loop configurations (open, closed, hybrid).

The thalamocortical circuitry of primates and rodents showed key differences, particularly regarding the presence of interneurons. Differences in interneuron involvement across species contribute to variations in circuit stability and responsiveness (Povysheva et al., 2006). Primates exhibit a more differentiated core and matrix structure in thalamic projections, with prominent interneuron-mediated inhibition, enhancing feedback loops and regulating spiking dynamics (Arcelli et al., 1997; A. Destexhe et al., 1998; Ilinsky et al., 1985; Rubio-Garrido et al., 2009; Yazdanbakhsh et al., 2023). Rodent models, on the other hand, provide valuable insights into closed-loop configurations, where their circuitry demonstrates heightened sensitivity to changes in thalamocortical strength (Moreira et al., 2025). Lacking significant interneuron activity (except in the LGN and few other first-order nuclei), rodent circuits rely more on direct thalamocortical excitation and inhibition, making them more sensitive to changes in thalamocortical connectivity (Cruikshank et al., 2010; Evangelio et al., 2018; O’Reilly et al., 2021; Rubio-Garrido et al., 2009; Simko & Markram, 2021). This sensitivity makes rodents an excellent model organism for studying fundamental principles of circuit plasticity and parameter-driven variations in synchrony. Additionally, the lack of significant interneuronal activity in rodent circuits allows for a more targeted examination of core network dynamics. These differences highlight the need for computational models to incorporate the distinct roles of interneurons and their dynamics to accurately simulate primate neurophysiology.

### Synchrony and Loop Configurations

In the closed-loop configurations, primate circuits consistently demonstrated higher synchrony, suggesting greater stability and resilience, possibly due to their more complex cortical mixing and connectivity patterns (Bhattacharya et al., 2021; Magrou et al., 2024). By contrast, the rodent closed loop exhibited reduced synchrony and greater sensitivity to changes in core thalamocortical strength. These findings emphasize the difference in roles between core and matrix projections in species with different cortical architectures.

The open loop highlighted significant differences in how rodent and primate thalamocortical circuits handle traveling waves. Rodent models exhibited persistent traveling waves across parameter variations, reflecting a fundamental instability in their loop configuration. Meanwhile, primate models demonstrated reduced traveling wave magnitude, reflecting more localized and efficient neural processing (Magrou et al., 2024).

The hybrid loop exhibited pronounced differences in synchrony dynamics. Rodents showed increased asymmetry and more pronounced traveling wave separations with increasing matrix strength and with greater recruitment of neurons. In contrast, primates demonstrated broader activity spread across cortical regions with increases in core, matrix, and cortico-cortical strength, without the pronounced asymmetry observed in rodents.

The spatiotemporal pattern of activity in the rodent model was more sensitive to changes in thalamocortical connectivity and exhibited greater variability across loop configurations compared to the primate model. This variability reflects the heightened sensitivity of rodent circuits to parameter changes, possibly due to their simpler loop architecture and fewer degrees of freedom compared to primates (Cruikshank et al., 2010; Laramée & Boire, 2015; O’Reilly et al., 2021). Parameter changes refer to adjustments in thalamocortical connectivity, including the strength of core and matrix projections and the degree of cortico-cortical mixing implemented in the model. While this sensitivity makes rodents an excellent model for studying fundamental mechanisms of network plasticity and parameter-driven dynamics, the richer thalamocortical loops observed in primates, with more degrees of freedom, provide greater stability and adaptability under varying conditions.

### Synchrony and Propagation Signatures Reveal Divergent TC Dynamics Across Species

The PCA and MPT analyses add a mechanistic interpretation to the qualitative differences observed across loop configurations. In primates, activity band angles remained tightly clustered near vertical (90°) across closed, open, and hybrid architectures, and MPT values remained consistently low, demonstrating that primate thalamocortical activity bands exhibit highly synchronous neuronal recruitment. Primate synchrony remained stable even when direct TRN feedback was removed (open loop) and connectivity parameter variations were pushed toward extreme values, highlighting the stability of primate synchrony. Rodent circuits behaved fundamentally differently. In multiple configurations, especially the closed loop and the open loop configurations, activity band angles deviated strongly from vertical, with values as low as 66° or exceeding 100°, and elevated MPT values up to ∼20 ms/neuron. These metrics reflect traveling-wave–like propagation and a substantial temporal spread across neurons, even when the corresponding primate condition remained synchronous. Only under specific connectivity parameter combinations, such as reduced core TC strength in the closed loop or increased core TC strength and higher cortico-cortical mixing in the hybrid loop, did rodent loop configurations approach the more synchronous patterns typically seen in primates. Even so, rodent synchrony remained less stable. Likewise, primate loops, while generally highly synchronous, displayed some desynchronization in cases where core TC strength or matrix TC strength was reduced within the closed and hybrid loops. Together, these results suggest that primate loops tend to remain synchronous across most configurations, while rodent loops showed a broader range of temporal patterns and reached synchrony only under specific connectivity conditions. In this context, the apparent stability of synchrony in primate circuits should not be interpreted as reduced circuit flexibility, but rather as preservation of synchronous population across a wide range of connectivity parameters. Importantly, synchrony represents only one dimension of network dynamics and does not fully capture functional flexibility. While rodent circuits exhibited flexibility through shifts between synchronous and traveling-wave dynamics, primates may preserve stable population synchrony while flexibly adjusting functional connectivity.

This divergence has important implications for interpreting species differences in spindle generation, sensory gating, and corticothalamic timing. Primate circuits may be tuned for more precise temporal coordination, while rodent circuits may be more prone to sequential or wave-like patterns that could be suited for different information processing.

### Common and Divergent Sleep Spindle Metrics in Rodents and Primates

Sleep spindles arise from dynamic thalamocortical interactions, with spindle variability reflecting distinct network states (Bazhenov et al., 2002). In our simulations, the raster of spindles and power spectra revealed clear species differences in how these network dynamics manifested (Figures 5 and 6). Across all loop configurations, the rodent spindle patterns appeared more irregular and heterogeneous across core and matrix than those in the primate model. In the raster plots, rodent spindles showed more variation in the shape and width, as well as greater inconsistency in how many neurons participated from spindle to spindle. In contrast, the primate simulations displayed spindles that were more uniform, with more consistent spindle shapes and clearer boundaries across core and matrix.

In terms of frequency, this same difference in regularity was reflected in the structure of the power spectra (Figure 6). Rodent simulations consistently showed flatter or broader frequency bands rather than sharply defined peaks, indicating more variable spindle frequencies. In contrast, the primate spectra tended to display clearer and more sharply defined power peaks, reflecting more uniform spindle frequencies, particularly in the hybrid core pathway, and showed less scattered power at neighboring frequencies. In rodents, multiple smaller peaks often appeared between 11 and 14 Hz, further suggesting a less uniform rhythmic organization of spindle activity.

Taken together, observing both the raster plots and the power spectra indicate that rodent spindle activity is more variable and less consistent, whereas primate spindles are more coherent and rhythmically uniform.

### Divergent Spindle-Band Signatures Across Rodents and Primates

The power spectrum of thalamocortical activity provided another source of insights regarding species-specific dynamics. “Fingerprint regions” in primates and rodents were revealed through analysis of average power versus frequency. For instance, primates exhibited characteristic shoulders in the spectrum, pointing to unique reliance on matrix-dominated pathways. Recent work identified statistically nonsignificant “frequency bumps” in rodents (Fernandez & Lüthi (2020). This variability underscores the need for further targeted analysis. Comparing primate and rodent models through individual power-frequency plots may provide greater clarity on these species-specific signatures.

Spectral analysis of thalamocortical activity revealed distinct frequency signatures in rodents and primates. In primates, EEG “fingerprint” regions, which are specific frequency bands associated with functional cortical specialization, were observed. Rodents, however, exhibited a different pattern, characterized by low-frequency oscillatory “shoulders” in the local field potential (LFP), indicating different modes of thalamocortical synchrony regulation. Further research utilizing EEG/MEG metrics could help determine how these spectral differences relate to cognitive processing and disease states.

Our results also revealed species-specific differences in thalamocortical spindle-band dynamics shaped by loop architecture. In open loop conditions, rodents and primates displayed similar general spectral power patterns; however, primates exhibited slightly more defined activity in the 13–14 Hz range, which was not seen in rodents (Figure 6A,D).

In closed loop conditions, rodents again exhibited activity in the 13–14 Hz range, but this power was diffuse and lacked a clear peak (Figure 6G–H). The most sharply defined spindle-band expression occurred in the lower-frequency range: the closed matrix configuration produced a narrow, high-amplitude peak at 11–12 Hz (Figure 6H), while the closed core loop showed a broader elevation spanning 10–12 Hz (Figure 6G). In contrast, primates showed only slight increases in power within the 13–14 Hz range and still lacked any distinct peaks in the 10–12 Hz band (Figure 6E–F). This absence of lower-frequency peaks was evident even in the closed matrix configuration, highlighting a divergence from the rodent profile.

Overall, rodents showed defined peaks in the lower spindle range under closed loop conditions, particularly at 11–12 Hz in the matrix pathway, whereas primates displayed more diffuse spectral power with increases in the 13–14 Hz range.

In hybrid loop configurations, species differences were most pronounced. Rodents displayed moderate activity between 13 and 14 Hz, though this remained relatively flat (Figure 6K–L). Several smaller peaks appeared across 11–14 Hz, with persistent elevation extending into 16 Hz, but no sharply localized feature emerged. In primates, however, hybrid core circuits revealed a clearly defined peak centered between 13 and 14 Hz, more pronounced than in any rodent condition (Figure 6I). A smaller, less distinct rise near 11 Hz was also occasionally observed (Figure 6J), suggesting the presence of a weaker alpha-range component. Overall, hybrid loops revealed the most distinct species differences, with primates showing a sharp and localized peak at 13–14 Hz, especially in the core circuit, while rodents exhibited flatter, more irregular power distributions across the same range.

Together, these findings point to a fundamental species difference: rodents exhibited strong, sharply localized activity in the lower spindle/alpha range (10–12 Hz) in both core and matrix closed loops (Figure 6G–H), while primates exhibited sharper spindle-band peaks centered at 13–14 Hz, particularly in the hybrid core configuration (Figure 6I). This divergence underscores distinct thalamocortical architectures shaping oscillatory dynamics across species.

Based on spectral data from rodent EEG recordings (Fernandez & Lüthi, 2020), sigma power tends to be low and lacks a distinct peak in the spindle frequency range. In contrast, our model’s pure rodent core loop configuration produced high-amplitude spindle activity (Figure 6G). This suggests that the rodent thalamocortical system may operate through a more integrated architecture, where core and matrix circuits blend and are not as segregated as they are in primates. This interpretation aligns with anatomical findings of mixed core and matrix TC connectivity in (Clascá et al., 2012; Rodriguez-Moreno et al., 2020; Rubio-Teves et al., 2024) and supports the need to incorporate matrix influence when modeling rodent thalamocortical dynamics. The rodent matrix loop in our model exhibited more diffuse spindle power, aligning with empirical observations of matrix-like or other blended circuit activity in rodents, which further highlights the importance of incorporating matrix-like circuit influence into rodent models of spindle dynamics. This pattern resembles, in part, the rodent power spectra reported in (Fernandez & Lüthi, 2020), which show less prominent spectral peaks (see their Figure 2). In contrast, simulations relying solely on a core-driven circuit tend to overestimate spindle power relative to *in vivo* recordings. These results suggest rodent thalamocortical circuits may need to be modeled as a functional hybrid, rather than as compartmentalized core and matrix subsystems.

In contrast, the primate thalamocortical loop demonstrates clearer structural and functional specialization between core and matrix circuits (Zikopoulos & Barbas, 2007b). In our model, the primate core loop produced two sharp peaks in the 13–15 Hz fast spindle range, particularly in the closed loop (Figure 2B), similar to the spectral patterns reported in Figure 2 of Fernandez & Lüthi (2020) which reported double peaks in somatosensory and auditory cortices and therefore are core-dominated. Our findings suggest that a substantial portion of closed-loop circuitry is present in these cortices, which are sensory, given that in our model the closed core configuration generated the two peaks. Matrix pathways, by comparison, yielded flatter, more spatially diffuse power profiles (Figure 6F,J), however, we observed a double-peak structure in hybrid matrix (Figure 6J) too, which carries the influence of core. Together, these may correspond to fast and slow spindle activity observed in EEG recordings, with our model suggesting that the EEG signal may be dominated by a combination of closed-core and hybrid-matrix activity.

In hybrid loop configurations, primate circuits displayed more structured spindle activity than rodents, with the hybrid matrix condition (Figure 6J) showing a distinct double-peak structure that may correspond to fast and slow spindle components. By contrast rodent hybrids exhibited flatter power profiles, particularly in both core and matrix hybrid configuration (Figure 6K–L), suggesting less distinct fast and slow spindle separation in rodents. This contrast further emphasizes how structural distinctions between species shape spindle expression across loop configurations.

These observations align with prior studies showing that human EEG typically displays dominant fast spindles, while rodent recordings, particularly from LFP of primary sensory areas (e.g., auditory or somatosensory cortex), often reflect lower-frequency components. The apparent flattening of the rodent spectra may be due to spatial convergence of core and matrix thalamocortical inputs across adjacent cortical layers (Clascá et al., 2012; Rodriguez-Moreno et al., 2020; Rubio-Teves et al., 2024). Although core and matrix neurons typically target distinct cortical layers (L4 and L1–3a, respectively), their projections in rodents often converge across adjacent laminae, resulting in spatial blending at the population level (Clascá et al., 2012; Rodriguez-Moreno et al., 2020; Rubio-Teves et al., 2024). When such input is averaged in silico, this convergence can reduce the sharpness of observed spectral peaks.

Although we compare our in-silico findings to previous empirical studies, it is important to note that the data sources differ in recording techniques. Prior work has primarily relied on EEG and local field potential (LFP) recordings, which capture activity at different spatial and temporal scales. Our model uses a basic averaging proxy across neuronal populations to approximate either LFP or EEG signals. This constitutes a gap between our model and the biological signals taken from EEG and LFP, as reported in Fernandez & Lüthi (2020), and our comparisons should be interpreted with that limitation in mind.

### Expanding the Role of the Cortex and Cortical Lamination in Thalamocortical Models

Beyond the interneurons and core-matrix dynamics, species differences in cortical expansion and laminar organization play a crucial role in shaping thalamocortical interactions. Primate cortices exhibit greater differentiation in cytoarchitecture which facilitates more complex cortico-cortical and cortico-thalamic feedback loops (Barbas & Zikopoulos, 2025; García-Cabezas et al., 2022b; Sherman & Guillery, 2002; Zikopoulos & Barbas, 2007b). This expanded cortical hierarchy allows for more refined top-down regulation of thalamic activity, influencing synchrony and oscillation dynamics. Rodent cortices, on the other hand, have more compact and less stratified laminar structure, leading to differences in thalamocortical connectivity patterns. The relatively simpler cortical organization in rodents likely results in more direct thalamic projections with less cortical feedback (Cappe et al., 2009; Ishizu et al., 2021), making these circuits more sensitive to parameter-driven fluctuations in spindles and synchrony. Additionally, the limited presence of a well-developed granular level (L4) in certain cortical levels in rodents may affect thalamic transmission, especially in sensory processing (Harris & Shepherd, 2015).

### Implications for Modeling of Neurodevelopmental Disorders

Based on our findings, *in silico* models of the thalamocortical loop should incorporate interneurons and gradient-based associative structures to more accurately reflect species-specific dynamics. Interneuronal activity, which is prominent in primates but significantly lower in rodent thalamus, plays a crucial role in shaping spiking activity and network dynamics. These distinctions must be integrated into computational models to ensure realistic simulations of thalamocortical function. Furthermore, rodent models may exhibit a stronger influence of matrix-dominated pathways, potentially impacting thalamocortical circuit stability and power-frequency relationships.

Additionally, observed differences in thalamocortical synchrony and spindle dynamics between rodents and primates provide a valuable framework for understanding the role of thalamocortical dysfunction in neurodevelopmental and neuropsychiatric disorders, such as autism spectrum disorder (ASD) and schizophrenia (SCZ) (Fernandez & Lüthi, 2020; Ferrarelli et al., 2010). Differences in core-matrix dynamics and synchrony suggest that computational models must integrate mechanisms that account for gradual, spatially structured changes in connectivity and synaptic strength to capture these pathophysiological features. Future studies could utilize EEG or MEG metrics to validate these findings and refine disorder-specific models.

## Supplementary Tables

**Supplementary Table S1.**
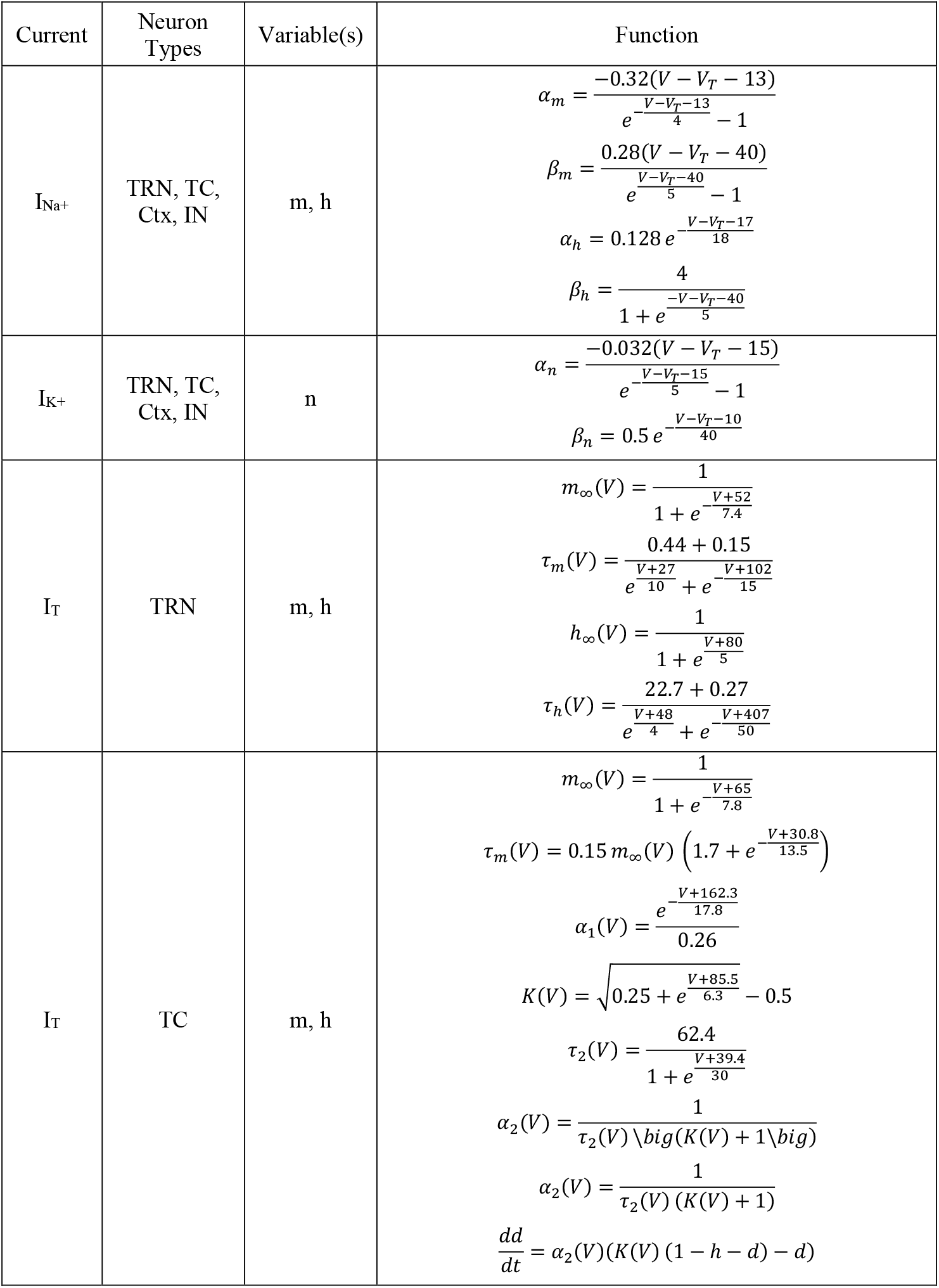
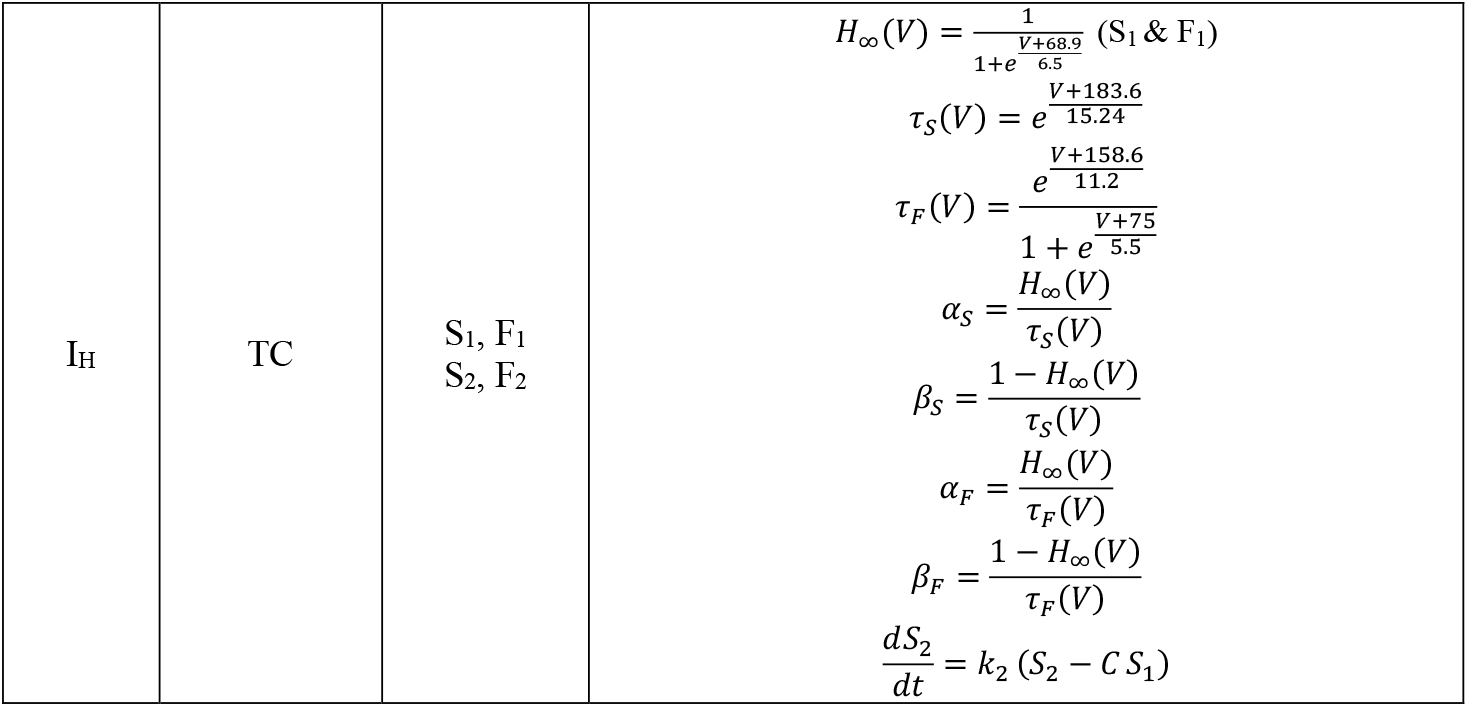
Voltage-gated and Intrinsic Currents Functions. The gating and time-constant functions are listed for each current, with the corresponding neuron types in which they are expressed: I_Na+_ (fast sodium), I_K+_ (delayed rectifier potassium), I_T_ (T-type calcium), I_H_ (hyperpolarization-activated current).

**Supplementary Table S2.** Neuron-Type Specific Constants. The maximal conductances 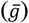, reversal potentials (E_i_), and initial voltages (V_initial_) are listed for Cortex (Ctx), thalamic reticular nucleus (TRN), thalamocortical relay nurons (TC), and interneurons (IN).

| Neuron Type | Currents | $\bar{g}$<br>mS/cm<sup>2</sup> | $E_i$<br>(mV) | $V_{initial}$ (mV) |
| --- | --- | --- | --- | --- |
| Cortex | $I_{Na}$ | 50 | 50 | -65 |
| | $I_K$ | 5 | -90 | |
| | $I_L$ | 0.1 | -70 | |
| TRN | $I_{Na}$ | 100 | 50 | -84 |
| | $I_K$ | 10 | -90 | |
| | $I_L$ | 0.05 | -75 | |
| | $I_T$ | 1.75 | 120 | |
| TC | $I_{Na}$ | 100 | 55 | -80 |
| | $I_K$ | 10 | -100 | |
| | $I_L$ | 50 | -86 | |
| | $I_H$ | 0.02 | -43 | |
| IN | $I_{Na}$ | 100 | 55 | -65 |
| | $I_K$ | 10 | -100 | |
| | $I_L$ | 50 | -86 | |

**Supplementary Table S3.** Ionotropic Synaptic Current Parameter Values. The forward (α) and backward (β) binding rates, and reversal potentials (E_j_), are listed for AMPA and GABA_A_ receptors.

| Current | $\alpha$ M<sup>-1</sup>S<sup>-1</sup> | $\beta$ S<sup>-1</sup> | E<sub>i</sub> (mV) |
| --- | --- | --- | --- |
| AMPA | 3.1842 | 0.1429 | 0 |
| GABA<sub>A</sub> | 0.53 | 0.184 | -100 |

**Supplementary Table S4.** Metabotropic Synaptic Current Parameter Values. The rate constants for neurotransmitter binding (K_1_) and unbinding (K_2_), second-messenger sensitization (K_3_) and desensitization (K_4_), the dissociation constant (K_d_), and the reversal potentials (E_j_) are listed for MGluR and GABA_B_ are provided.

| Metabotropic Currents | K<sub>1</sub><br>M<sup>-1</sup>S<sup>-1</sup> | K<sub>2</sub><br>S<sup>-1</sup> | K<sub>3</sub><br>S<sup>-1</sup> | K<sub>4</sub><br>S<sup>-1</sup> | K<sub>d</sub><br>μM<sup>2</sup> | E<sub>i</sub><br>(mV) |
| --- | --- | --- | --- | --- | --- | --- |
| MGluR | 1.3 | 6.6 | 1.9 | 0.0064 | 100 | 0 |
| GABA<sub>B</sub> | 1.3 | 6.6 | 0.9 | 0.0064 | 100 | -100 |

## Bibliography

Arcaro, M. J., Pinsk, M. A., & Kastner, S. (2015). The Anatomical and Functional Organization of the Human Visual Pulvinar. Journal of Neuroscience, 35(27), 9848–9871. 10.1523/JNEUROSCI.1575-14.2015

Arcelli, P., Frassoni, C., Regondi, M. C., Biasi, S. D. E., & Spreafico, R. (1997). GABAergic Neurons in Mammalian Thalamus: A Marker of Thalamic Complexity? Brain Research Bulletin, 42(1), 27–37. 10.1016/S0361-9230(96)00107-4

Bal, T., & McCormick, D. A. (1993). Mechanisms of oscillatory activity in guinea-pig nucleus reticularis thalami in vitro: a mammalian pacemaker. The Journal of Physiology, 468(1), 669–691. 10.1113/jphysiol.1993.sp019794

Barbas, H., & Zikopoulos, B. (2025). The Cortical Structural Model Extends to Thalamocortical Connections. European Journal of Neuroscience, 61(12). 10.1111/ejn.70167

Bazhenov, M., Timofeev, I., Steriade, M., & Sejnowski, T. J. (2002). Model of Thalamocortical Slow-Wave Sleep Oscillations and Transitions to Activated States. The Journal of Neuroscience, 22(19), 8691–8704. 10.1523/JNEUROSCI.22-19-08691.2002

Bhattacharya, S., Cauchois, M. B. L., Iglesias, P. A., & Chen, Z. S. (2021). The impact of a closed-loop thalamocortical model on the spatiotemporal dynamics of cortical and thalamic traveling waves. Scientific Reports, 11(1), 14359. 10.1038/s41598-021-93618-6

Brown, J. W., Taheri, A., Kenyon, R. V, Berger-Wolf, T. Y., & Llano, D. A. (2020). Signal Propagation via Open-Loop Intrathalamic Architectures: A Computational Model. Eneuro, 7(1), ENEURO.0441-19.2020. 10.1523/ENEURO.0441-19.2020

Cappe, C., Morel, A., Barone, P., & Rouiller, E. M. (2009). The Thalamocortical Projection Systems in Primate: An Anatomical Support for Multisensory and Sensorimotor Interplay. Cerebral Cortex, 19(9), 2025–2037. 10.1093/cercor/bhn228

Castelnovo, A., D’Agostino, A., Mayeli, A., Albantakis, L., Tononi, G., & Ferrarelli, F. (2025). Sleep Spindle Abnormalities as Neurophysiological Biomarkers of Schizophrenia Spectrum Disorders: From Cellular Mechanisms and Neural Circuits to Clinical Implications. Biological Psychiatry, 98(11), 809–818. 10.1016/j.biopsych.2025.06.014

Chartrand, T., Dalley, R., Close, J., Goriounova, N. A., Lee, B. R., Mann, R., Miller, J. A., Molnar, G., Mukora, A., Alfiler, L., Baker, K., Bakken, T. E., Berg, J., Bertagnolli, D., Braun, T., Brouner, K., Casper, T., Csajbok, E. A., Dee, N., … Lein, E. S. (2023). Morphoelectric and transcriptomic divergence of the layer 1 interneuron repertoire in human versus mouse neocortex. Science, 382(6667). 10.1126/science.adf0805

Clascá, F., Rubio-Garrido, P., & Jabaudon, D. (2012). Unveiling the diversity of thalamocortical neuron subtypes. European Journal of Neuroscience, 35(10), 1524–1532. 10.1111/j.1460-9568.2012.08033.x

Cruikshank, S. J., Urabe, H., Nurmikko, A. V, & Connors, B. W. (2010). Pathway-Specific Feedforward Circuits between Thalamus and Neocortex Revealed by Selective Optical Stimulation of Axons. Neuron, 65(2), 230–245. 10.1016/j.neuron.2009.12.025

Destexhe, a, Mainen, Z. F., & Sejnowski, T. J. (1998). Kinetic models of synaptic transmission: From Ions to Networks. Methods in Neural Modeling: From Ions to Networks, 1–25.

Destexhe, A., Babloyantz, A., & Sejnowski, T. J. (1993). Ionic mechanisms for intrinsic slow oscillations in thalamic relay neurons. Biophysical Journal, 65(4), 1538–1552. 10.1016/S0006-3495(93)81190-1

Destexhe, A., Contreras, D., Sejnowski, T. J., & Steriade, M. (1994). A model of spindle rhythmicity in the isolated thalamic reticular nucleus. Journal of Neurophysiology, 72(2), 803–818. 10.1152/jn.1994.72.2.803

Destexhe, A., Contreras, D., & Steriade, M. (1998). Mechanisms underlying the synchronizing action of corticothalamic feedback through inhibition of thalamic relay cells. Journal of Neurophysiology, 79(2), 999–1016. 10.1152/jn.1998.79.2.999

Destexhe, A., Contreras, D., Steriade, M., Sejnowski, T. J., & Huguenard, J. R. (1996). In vivo, in vitro, and computational analysis of dendritic calcium currents in thalamic reticular neurons. The Journal of Neuroscience, 16(1), 169–185. 10.1523/JNEUROSCI.16-01-00169.1996

Evangelio, M., García-Amado, M., & Clascá, F. (2018). Thalamocortical Projection Neuron and Interneuron Numbers in the Visual Thalamic Nuclei of the Adult C57BL/6 Mouse. Frontiers in Neuroanatomy, 12. 10.3389/fnana.2018.00027

Fernandez, L. M. J., & Lüthi, A. (2020). Sleep Spindles: Mechanisms and Functions. Physiological Reviews, 100(2), 805–868. 10.1152/physrev.00042.2018

Ferrarelli, F., Peterson, M. J., Sarasso, S., Riedner, B. A., Murphy, M. J., Benca, R. M., Bria, P., Kalin, N. H., & Tononi, G. (2010). Thalam ic Dysfunction in Schizophrenia Suggested by Whole-Night Defi cits in Slow and Fast Spindles. 1339–1348.

García-Cabezas, M.Á., Hacker, J. L., & Zikopoulos, B. (2022a). Homology of neocortical areas in rats and primates based on cortical type analysis: an update of the Hypothesis on the Dual Origin of the Neocortex. Brain Structure and Function, 228(5), 1069–1093. 10.1007/s00429-022-02548-0

García-Cabezas, M.Á., Hacker, J. L., & Zikopoulos, B. (2022b). Homology of neocortical areas in rats and primates based on cortical type analysis: an update of the Hypothesis on the Dual Origin of the Neocortex. Brain Structure and Function, 228(5), 1069–1093. 10.1007/s00429-022-02548-0

Gerardo, C.-M., & Manuel, M.-M. V. (2020). The thalamic reticular nucleus: A common nucleus of neuropsychiatric diseases and deep brain stimulation. Journal of Clinical Neuroscience, 73, 1–7. 10.1016/j.jocn.2020.01.061

Gonzalez, C., Jiang, X., Gonzalez-Martinez, J., & Halgren, E. (2022). Human Spindle Variability. The Journal of Neuroscience, 42(22), 4517–4537. 10.1523/jneurosci.1786-21.2022

Harris, K. D., & Shepherd, G. M. G. (2015). The neocortical circuit: themes and variations. Nature Neuroscience, 18(2), 170–181. 10.1038/nn.3917

Hodgkin, A. L., & Huxley, A. F. (1952). A quantitative description of membrane current and its application to conduction and excitation in nerve. The Journal of Physiology, 117(4), 500–544. 10.1113/jphysiol.1952.sp004764

Huguenard, J. R., & Prince, D. A. (1992). A novel T-type current underlies prolonged Ca(2+)-dependent burst firing in GABAergic neurons of rat thalamic reticular nucleus. The Journal of Neuroscience, 12(10), 3804–3817. 10.1523/JNEUROSCI.12-10-03804.1992

Ilinsky, I. A., Jouandet, M. L., & Goldman-Rakic, P. S. (1985). Organization of the nigrothalamocortical system in the rhesus monkey. Journal of Comparative Neurology, 236(3), 315–330. 10.1002/cne.902360304

Ishizu, K., Shiramatsu, T. I., Hitsuyu, R., Oizumi, M., Tsuchiya, N., & Takahashi, H. (2021). Information flow in the rat thalamo-cortical system: spontaneous vs. stimulus-evoked activities. Scientific Reports, 11(1), 19252. 10.1038/s41598-021-98660-y

Jones, E. G. (1998). Commentary: ViewPoint: The Core and Matrix of Thalamic Organization. 2, 331– 345.

Jones, E. G. (2007). The Thalamus (Second). Cambridge University Press.

Jorstad, N. L., Song, J. H. T., Exposito-Alonso, D., Suresh, H., Castro-Pacheco, N., Krienen, F. M., Yanny, A. M., Close, J., Gelfand, E., Long, B., Seeman, S. C., Travaglini, K. J., Basu, S., Beaudin, M., Bertagnolli, D., Crow, M., Ding, S.-L., Eggermont, J., Glandon, A., … Bakken, T. E. (2023). Comparative transcriptomics reveals human-specific cortical features. Science, 382(6667). 10.1126/science.ade9516

Joyce, M. K. P., Marshall, L. G., Banik, S. L., Wang, J., Xiao, D., Bunce, J. G., & Barbas, H. (2022). Pathways for Memory, Cognition and Emotional Context: Hippocampal, Subgenual Area 25, and Amygdalar Axons Show Unique Interactions in the Primate Thalamic Reuniens Nucleus. The Journal of Neuroscience, 42(6), 1068–1089. 10.1523/JNEUROSCI.1724-21.2021

Kim, C. N., Shin, D., Wang, A., & Nowakowski, T. J. (2023). Spatiotemporal molecular dynamics of the developing human thalamus. Science, 382(6667). 10.1126/science.adf9941

Laramée, M.-E., & Boire, D. (2015). Visual cortical areas of the mouse: comparison of parcellation and network structure with primates. Frontiers in Neural Circuits, 8. 10.3389/fncir.2014.00149

Latchoumane, C.-F. V., Ngo, H.-V. V., Born, J., & Shin, H.-S. (2017). Thalamic Spindles Promote Memory Formation during Sleep through Triple Phase-Locking of Cortical, Thalamic, and Hippocampal Rhythms. Neuron, 95(2), 424-435.e6. 10.1016/j.neuron.2017.06.025

Magrou, L., Joyce, M. K. P., Froudist-Walsh, S., Datta, D., Wang, X.-J., Martinez-Trujillo, J., & Arnsten, A. F. T. (2024). The meso-connectomes of mouse, marmoset, and macaque: network organization and the emergence of higher cognition. Cerebral Cortex, 34(5), bhae174. 10.1093/cercor/bhae174

Manoach, D. S., & Stickgold, R. (2019a). Abnormal Sleep Spindles, Memory Consolidation, and Schizophrenia. Annual Review of Clinical Psychology, 15(1), 451–479. 10.1146/annurev-clinpsy-050718-095754

Manoach, D. S., & Stickgold, R. (2019b). Abnormal Sleep Spindles, Memory Consolidation, and Schizophrenia. Annual Review of Clinical Psychology, 15, 451–479. 10.1146/annurev-clinpsy-050718-095754

McCormick, D. A., & Huguenard, J. R. (1992). A model of the electrophysiological properties of thalamocortical relay neurons. Journal of Neurophysiology, 68(4), 1384–1400. 10.1152/jn.1992.68.4.1384

Mengxing, L., Lerma-Usabiaga, G., Clascá, F., & Paz-Alonso, P. M. (2023). High-Resolution Tractography Protocol to Investigate the Pathways between Human Mediodorsal Thalamic Nucleus and Prefrontal Cortex. The Journal of Neuroscience, 43(46), 7780–7798. 10.1523/JNEUROSCI.0721-23.2023

Moreira, J. V., Borges, F. S., Atherton, Z., Crandall, S. R., Varela, C., & Dura-Bernal, S. (2025). Closed-Loop Connectivity Best Supports Angular Tuning and Sleep Dynamics in a Biophysical Thalamocortical Circuit Model. 10.1101/2025.08.18.670921

Murray, K. D., Choudary, P. V., & Jones, E. G. (2007). Nucleus- and cell-specific gene expression in monkey thalamus. Proceedings of the National Academy of Sciences, 104(6), 1989–1994. 10.1073/pnas.0610742104

Mylonas, D., Machado, S., Larson, O., Patel, R., Cox, R., Vangel, M., Maski, K., Stickgold, R., & Manoach, D. S. (2022). Dyscoordination of non-rapid eye movement sleep oscillations in autism spectrum disorder. Sleep, 45(3). 10.1093/sleep/zsac010

O’Reilly, C., Iavarone, E., Yi, J., & Hill, S. L. (2021). Rodent somatosensory thalamocortical circuitry: Neurons, synapses, and connectivity. Neuroscience & Biobehavioral Reviews, 126, 213–235. 10.1016/j.neubiorev.2021.03.015

Pinault, D. (2004). The thalamic reticular nucleus: structure, function and concept. Brain Research Reviews, 46(1), 1–31. 10.1016/j.brainresrev.2004.04.008

Pospischil, M., Toledo-Rodriguez, M., Monier, C., Piwkowska, Z., Bal, T., Frégnac, Y., Markram, H., & Destexhe, A. (2008). Minimal Hodgkin-Huxley type models for different classes of cortical and thalamic neurons. Biological Cybernetics, 99(4–5), 427–441. 10.1007/s00422-008-0263-8

Povysheva, N. V, Gonzalez-Burgos, G., Zaitsev, A. V, Kröner, S., Barrionuevo, G., Lewis, D. A., & Krimer, L. S. (2006). Properties of Excitatory Synaptic Responses in Fast-spiking Interneurons and Pyramidal Cells from Monkey and Rat Prefrontal Cortex. Cerebral Cortex, 16(4), 541–552. 10.1093/cercor/bhj002

Reichova, I., & Sherman, S. M. (2004). Somatosensory Corticothalamic Projections: Distinguishing Drivers From Modulators. Journal of Neurophysiology, 92(4), 2185–2197. 10.1152/jn.00322.2004

Rodriguez-Moreno, J., Porrero, C., Rollenhagen, A., Rubio-Teves, M., Casas-Torremocha, D., Alonso-Nanclares, L., Yakoubi, R., Santuy, A., Merchan-Pérez, A., DeFelipe, J., Lübke, J. H. R., & Clasca, F. (2020). Area-Specific Synapse Structure in Branched Posterior Nucleus Axons Reveals a New Level of Complexity in Thalamocortical Networks. The Journal of Neuroscience, 40(13), 2663– 2679. 10.1523/JNEUROSCI.2886-19.2020

Rubio-Garrido, P., Pérez-de-Manzo, F., Porrero, C., Galazo, M. J., & Clascá, F. (2009). Thalamic Input to Distal Apical Dendrites in Neocortical Layer 1 Is Massive and Highly Convergent. Cerebral Cortex, 19(10), 2380–2395. 10.1093/cercor/bhn259

Rubio-Teves, M., Martín-Correa, P., Alonso-Martínez, C., Casas-Torremocha, D., García-Amado, M., Timonidis, N., Sheiban, F. J., Bakker, R., Tiesinga, P., Porrero, C., & Clascá, F. (2024). Beyond Barrels: Diverse Thalamocortical Projection Motifs in the Mouse Ventral Posterior Complex. The Journal of Neuroscience, 44(43), e1096242024. 10.1523/JNEUROSCI.1096-24.2024

Saalmann, Y. B., Pinsk, M. A., Wang, L., Li, X., & Kastner, S. (2012). The Pulvinar Regulates Information Transmission Between Cortical Areas Based on Attention Demands. Science, 337(6095), 753–756. 10.1126/science.1223082

Sherman, S. M., & Guillery, R. W. (2002). The role of the thalamus in the flow of information to the cortex. Philosophical Transactions of the Royal Society of London. Series B: Biological Sciences, 357(1428), 1695–1708. 10.1098/rstb.2002.1161

Simko, J., & Markram, H. (2021). Morphology, physiology and synaptic connectivity of local interneurons in the mouse somatosensory thalamus. The Journal of Physiology, 599(22), 5085– 5101. 10.1113/JP281711

Timbie, C., García-Cabezas, M.Á., Zikopoulos, B., & Barbas, H. (2020). Organization of primate amygdalar–thalamic pathways for emotions. PLOS Biology, 18(2), e3000639. 10.1371/journal.pbio.3000639

Whyte, C. J., Redinbaugh, M. J., Shine, J. M., & Saalmann, Y. B. (2024). Thalamic contributions to the state and contents of consciousness. Neuron, 112(10), 1611–1625. 10.1016/j.neuron.2024.04.019

Willis, A. M., Slater, B. J., Gribkova, E. D., & Llano, D. A. (2015). Open-loop organization of thalamic reticular nucleus and dorsal thalamus: a computational model. Journal of Neurophysiology, 114(4), 2353–2367. 10.1152/jn.00926.2014

Xiao, D., Zikopoulos, B., & Barbas, H. (2009). Laminar and modular organization of prefrontal projections to multiple thalamic nuclei. Neuroscience, 161(4), 1067–1081. 10.1016/j.neuroscience.2009.04.034

Yazdanbakhsh, A., Barbas, H., & Zikopoulos, B. (2023). Sleep spindles in primates: Modeling the effects of distinct laminar thalamocortical connectivity in core, matrix, and reticular thalamic circuits. Network Neuroscience, 7(2), 743–768. 10.1162/netn_a_00311

Zikopoulos, B., & Barbas, H. (2007a). Circuits formultisensory integration and attentional modulation through the prefrontal cortex and the thalamic reticular nucleus in primates. Reviews in the Neurosciences, 18(6), 417–438. 10.1515/revneuro.2007.18.6.417

Zikopoulos, B., & Barbas, H. (2007b). Parallel driving and modulatory pathways link the prefrontal cortex and thalamus. PLoS ONE, 2(9). 10.1371/journal.pone.0000848

